# Dysregulation of GABAergic System in the Hippocampus and Inferior Colliculi of Rats during the Development of Audiogenic Epilepsy

**DOI:** 10.1101/2023.04.20.537676

**Authors:** Andrey P. Ivlev, Elena V. Chernigovskaya, Svetlana D. Nikolaeva, Alexey A. Kulikov, Margarita V. Glazova, Alexandra A. Naumova

**Affiliations:** Sechenov Institute of Evolutionary Physiology and Biochemistry, the Russian Academy of Sciences, St. Petersburg, Russia

**Keywords:** Krushinsky-Molodkina rats, audiogenic seizures, inferior colliculi, hippocampus, postnatal ontogenesis

## Abstract

Epilepsy is tightly associated with dysfunction of inhibitory γ-aminobutyric acid (GABA) neurotransmission. In this study, Krushinsky-Molodkina (KM) rats genetically prone to audiogenic seizures (AGS) were used. KM rats are characterized by the development of audiogenic epilepsy during postnatal ontogenesis, with AGS onset at the age of 1.5-2 months and fully developed AGS expression by 3^rd^ month. We analyzed GABAergic system of the inferior colliculi (IC) and the hippocampus of KM rats at different stages of postnatal development. Wistar rats were used as a control. In the IC of young KM rats, Na^+^/K^+^/Cl^−^ cotransporter 1 (NKCC1) expression was increased, while in adults K^+^/Cl^−^ cotransporter 2 (KCC2) was downregulated indicating impairment of postsynaptic GABA action both at early and later stages of postnatal development. In the hippocampus of young KM rats, we observed a decrease in activity of GABAergic neurons and downregulation of KCC2 and NKCC1. Adult rats, in opposite, demonstrated elevated activity of the hippocampal GABAergic cells and unchanged expression of chlorine transporters indicating upregulation of GABA transmission. Thus, GABA dysregulation in the IC can mediate the seizure susceptibility in adult KM rats, while in the hippocampus, upregulation of GABAergic system can restrict the spreading of epileptiform activity from the brainstem.

## Introduction

Nowadays, a growing body of evidence indicates that impairment of inhibitory γ-aminobutyric acid (GABA) neurotransmission in the brain is one of the key mechanisms of epilepsy development [1]. Thus, the studies of clinical material from patients with epilepsy and experimental models have demonstrated loss of GABAergic neurons, attenuation of GABA production, and a decrease in GABA receptors on the membranes of GABA target cells [2-5]. Moreover, epileptiform activity can be associated with alterations in neuronal expression of Cl^−^ transporters – K^+^/Cl^−^ cotransporter 2 (KCC2) and Na^+^/K^+^/Cl^−^ cotransporter 1 (NKCC1) that lead to the impairment of Cl^−^ balance and, as a result, to attenuation of GABAergic inhibition and even switch of GABA effect from inhibition to excitation [6, 7].

On the other hand, epileptogenesis can be associated with altered genetic control of GABAergic system. It was shown that mutations in *GAD1* gene coding the crucial enzyme of GABA biosynthesis, glutamic acid decarboxylase 67 (GAD67), are associated with severe violations in the brain development including epileptic encephalopathy [8]. In patients with epilepsy, multiple mutations of GABA receptors were also detected [9], and genetic inactivation of KCC2 in mice resulted in increased seizure susceptibility [10]. Although such data are fragmentary and only partially characterize the causes of violations, they suggest that impaired functioning of GABAergic system can be not only the consequence of epileptiform activity, but also the cause of epilepsy development.

The perspective experimental approach to reveal genetically determined mechanisms of epileptogenesis is using of the animals with genetic predisposition to reflex (audiogenic) seizures. Nowadays, there are several well-studied strains of rats with audiogenic sensitivity including genetically epilepsy-prone rats (GEPR), Wistar audiogenic rats (WAR), Wistar audiogenic rats from Strasbourg (WAS), аnd Krushinsky - Molodkina (КМ) rats [11]. Increased seizure susceptibility develops in such animals during postnatal ontogenesis making these models especially convenient to investigate mechanisms and dynamics of the development of inherited epilepsy in humans.

Electrophysiological studies of animals with audiogenic sensitivity have shown that single stimulations of audiogenic seizures (AGS) induce epileptiform activity in the brainstem, and the key center responsible for AGS triggering is the inferior colliculi (IC) of the quadrigemina [12-14]. The hippocampus and other elements of the limbic system, apparently, are not involved in the expression of single AGS, being recruited in epileptic circuits only during repetitive AGS stimulations, so-called audiogenic kindling [15, 16]. At the same time, previous studies have shown that audiogenic animals demonstrate a number of alterations in the GABAergic system of both the IC and the hippocampus, including abnormally low GABA production and decreased KCC2 expression [17-20]. Indeed, such alterations could contribute to hyperexcitability of these brain structures and support the development of reflex epilepsy. However, GABA imbalance was mainly observed in sound-stimulated animals, making it difficult to distinguish inherited and seizure-induced changes. Nevertheless, it is reasonable to suppose that in audiogenic rats, postnatal development is associated with abnormal formation of neural circuits which is mediated by the presence of genetically determined aberrations in the GABAergic system and provides the basis for seizure susceptibility in adulthood.

In rodents, first postnatal weeks consist a critical period for the establishment of GABAergic system in the IC and hippocampus. In the IC, GABAergic cells are detected since the 8^th^ postnatal day (P8), and by the end of the first month of life their number no longer differs from adult animals [21]. In the hippocampus, the first GABAergic cells appear already during prenatal development (at 17^th^ – 18^th^ embryonic days, E17 – E18) [22] but only by P20 GABA production in the hippocampus reaches the adult level [23]. Moreover, early postnatal development is associated with a gradual increase in KCC2 expression in neurons which is necessary for the establishment of inhibitory (hyperpolarizing) GABA action [24, 25]. On the other hand, there are practically no data about the development of GABAergic system in laboratory rodents with hereditary epilepsy. To fill in, at least in part, the missing information, we have analyzed maturation of this system the hippocampus and IC of KM rats during postnatal ontogenesis.

Audiogenic reflex epilepsy in KM rats is completely formed by the 3^rd^ month of life [26]. At earlier stages of postnatal development, KM rats either do not respond to sound stimulation or demonstrate incomplete pattern of seizure that usually includes only wild running phase. Previously we demonstrated the delay in postnatal morphogenesis of both the IC and the hippocampus of audiogenic KM rats in comparison with control Wistar rats [27, 28]. Moreover, analysis of GAD67, parvalbumin (PV), and synapsin 1 revealed abnormal functioning of GABAergic neurons in the IC of KM rats both in young and adult state [29]. In the present study, we have continued investigation of the IC in KM rats with special focus on GABAergic transmission. Moreover, we performed a comprehensive analysis of GABAergic system postnatal development in the hippocampus of these animals. Obtained results let us to disclose genetically determined alterations in the maturation of inhibitory circuits in the IC and the hippocampus and suggest the contribution of observed changes into the establishment of increased convulsive readiness in audiogenic rodents.

## Results

In our experiments, we used naïve KM rats which had no experience of AGS expression. Wistar rats were used as a control. Animals of three age groups were investigated: 1) 15 days (P15) when active morphogenesis of the IC and the hippocampus occurs; 2) 2 month (P60) when the development of both structures of interest is completed, but KM rats do not express stable AGS; 3) 4 months (P120) when the ability of KM rats to demonstrate express AGS is completely formed.

### Analysis of GABAergic system in the inferior colliculi (IC)

Firstly, we analyzed expression of vesicular GABA transporter (VGAT) which is responsible for loading of GABA from cytoplasm of GABAergic neurons into synaptic vesicles [30]. Our data revealed no differences in VGAT expression in the IC between KM and Wistar rats at P15 and P120, however, in the IC of 2-months-old KM rats (P60) was significantly upregulated (Figure 1a, b).

**Figure 1.**
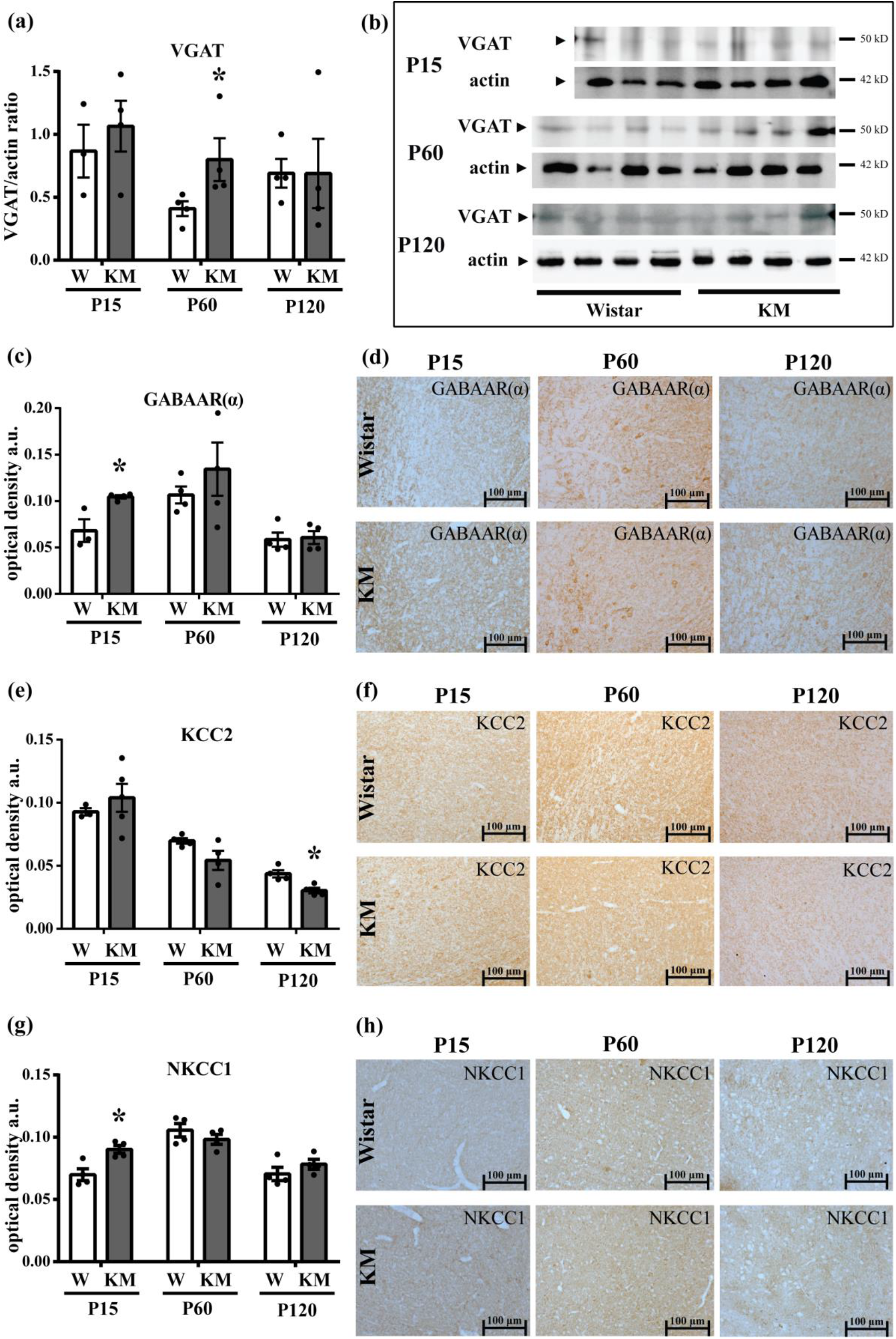
Expression of proteins responsible for γ-aminobutyric acid (GABA) transmission in the inferior colliculi (IC) of Krushinsky – Molodkina (KM) and Wistar rats during postnatal ontogenesis. **(a)** Western blot analysis of vesicular GABA transporter (VGAT) in the IC at 15^th^ (P15) and 120^th^ (P120) days of postnatal development did not reveal any differences between Wistar (W) and KM rats, however, at 60^th^ day (P60), VGAT expression in the IC of KM rats was higher than in the control. Western blot data are presented as mean ± SEM. * – p<0.05 vs control. **(b)** Representative immunoblot images of VGAT and actin in the hippocampus of KM and Wistar rats at corresponding ontogenetic stages. **(c)** Immunohistochemical assay demonstrated increased expression of α1-subunit of GABA-A receptors (GABAAR(α)) in the IC of KM rats at P15, while at other points it did not differ from Wistar rats. **(d)** Representative micrographs of GABAAR(α) immunostaining in the IC central nucleus of Wistar and KM rats at P15, P60, and P120. **(e)** Expression of K+/Cl^−^ cotransporter (KCC2) in the IC of KM rats did not differ from Wistar control at P15 and P60 but was significantly reduced at P120. **(f)** Representative micrographs of KCC2 immunostaining in the IC central nucleus of Wistar and KM rats at P15, P60, and P120. **(g)** Expression of Na+/K+/Cl^−^ cotransporter 1 (NKCC1) in the IC of KM rats, in opposite, was elevated at P15, but not at other time points. **(h)** Representative micrographs of NKCC1 immunostaining in the IC central nucleus of Wistar and KM rats at P15, P60, and P120. Plots **(c), (e), (g)** demonstrate optical density of immunopositive substance in arbitrary units (a.u.). Immunohistochemical data are shown as mean ± standard error of mean (SEM). * – p<0.05 vs control. Scale bars: 100 µm.

To estimate postsynaptic effects of GABA in the IC, we investigated expression of GABA-A receptor α1-subunit (GABAAR(α)), which is the obligatory component of the receptor participating in GABA binding [31]. Immunohistochemical analysis revealed increased expression of GABAAR(α) in KM rats on the 15^th^ day of life, while in other age groups (P60, P120) it did not differ from corresponding Wistar control (Figure 1c, d).

In addition, we analyzed the expression of active Cl^−^ transporters, KCC2 and NKCC1, which maintain Cl^−^ balance in GABA target cells by accumulating or extruding Cl^−^, respectively [6]. Our data showed that at P15 KCC2 expression in the IC of KM rats was unchanged (Figure 1e, f), while NKCC1 expression was higher than in Wistar rats, probably, indicating the increase in Cl^−^ influx (Figure 1g, h). At P60 expression of both transporters did not differ between KM and Wistar rats (Figure 1e-h). However, 4-month-old KM rats demonstrated abnormally low KCC2 expression along with unchanged NKCC1 (Figure 1e-h). Observed changes could point on altered intracellular Cl^−^ balance with up-regulation of its efflux in adult animals.

### Analysis of GABAergic system in the hippocampus

#### 15^th^ postnatal day (P15)

To analyze the activity of GABA synthesis in the hippocampus, we evaluated the expression of GAD67, the key marker of GABAergic interneurons [32]. In the hippocampus of 15-day-old KM rats, the expression of GAD67 mRNA was significantly lower than in Wistar rats (Figure 2a). Immunohistochemical assay showed ubiquitous decrease of GAD67 expression in the processes of GABAergic neurons (Figure 2c, d), while the numbers of GAD67-immunopositive cells were, in opposite, elevated in the hilus and cornu Ammonis 4 subfield (CA4), but not in the dentate gyrus (DG) granular layer (Figure 2b, d). This contradiction was obviously associated with impaired transport of the enzyme from the cell bodies to the axons.

**Figure 2.**
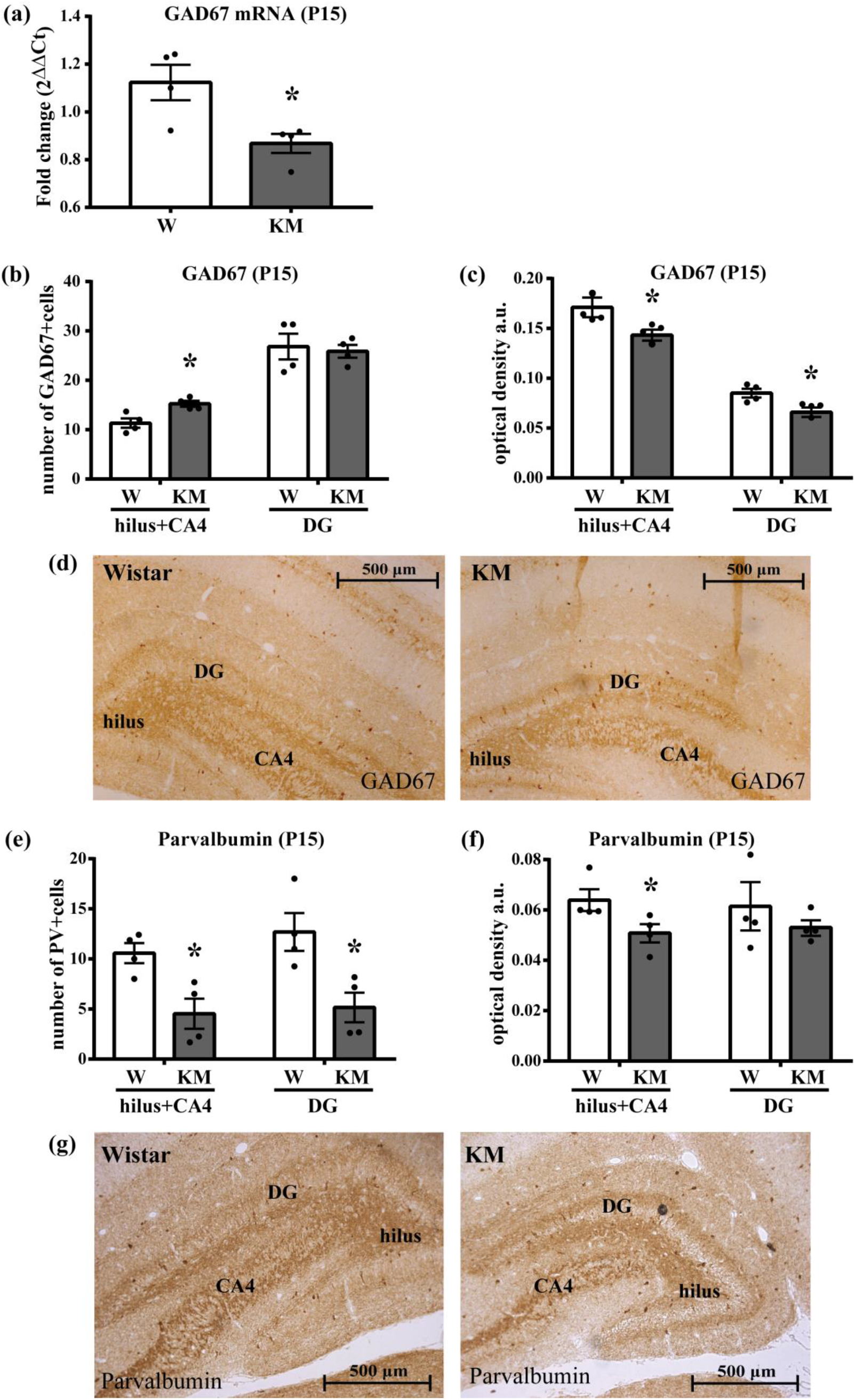
Analysis of glutamic acid decarboxylase 67 (GAD67) and parvalbumin (PV) in the hippocampus of 15-days-old Wistar and KM rats. **(a)** Real-time PCR analysis revealed significantly decreased GAD67 mRNA in the hippocampus of KM rats in comparison with Wistar (W) control. Data are presented as mean ± SEM. * – p<0.05 vs control. **(b)** Immunohistochemical assay detected increased number of GAD67-positive cells in the hilus and cornu Ammonis 4 subfield (hilus+CA4) of KM rats, however, in the dentate gyrus (DG) granular layer, no changes were observed. **(c)** GAD67 expression in GABAergic fibers was significantly decreased in both the (hilus+CA4) zone and DG granular layer of KM rats. **(d)** Representative micrographs of GAD67 immunostaining in the hippocampus of 15-day-old Wistar and KM rats. **(e, f)** KM rats demonstrated decreased numbers of PV-positive cells in both investigated hippocampal zones **(e)**, while PV expression in fibers **(f)** was lower in the hilus and CA4, but not in the DG granular layer. **(g)** Representative micrographs of parvalbumin immunostaining in the hippocampus of Wistar and KM rats at P15. Plots **(b)** and **(e)** demonstrate numbers of immunopositive cells; plots **(c)** and **(f)** – optical density of immunopositive substance, arbitrary units (a.u.). Immunohistochemical data are presented as mean ± SEM. * – p < 0.05 vs control. Scale bars: 500 µm.

Ca^2+^-binding protein parvalbumin (PV) is also widely used as a marker of GABAergic neurons [33, 34]. It participates in buffering of intracellular Ca^2+^ in cells with high electrical and metabolic activity. As PV-positive neurons form the most part of GABAergic synapses with projection neurons, this population of cells is considered to be the main component of GABAergic system in the hippocampus [33]. In 15-day-old KM rats, populations of PV-positive neurons were decreased in all investigated hippocampal regions (Figure 2e, g), while PV expression in GABAergic fibers was lower only in the hilus and CA4 (Figure 2f, g).

To estimate the synaptic transmission in the hippocampus, we analyzed expression and phosphorylation of synapsin 1. In unphosphorylated form, this protein participates in binding of synaptic vesicles to the cytoskeleton, while phosphorylated synapsin 1 controls vesicle release allowing neurotransmitter exocytosis [35]. Our results showed unchanged expression of synapsin 1 (Figure 3a, f) in the hippocampus of 15-days-old KM rats. At the same time decreased phosphorylation of synapsin 1 at Ser62/67 (Figure 3b, f) pointed on a decrease in its activity and downregulation of neurotransmitter release. These changes were accompanied with elevated expression of VGAT (Figure 3c, f) and GABAAR(α) (Figure 3e, f). However, expression of GABAAR(α) mRNA was decreased (Figure 3d).

**Figure 3.**
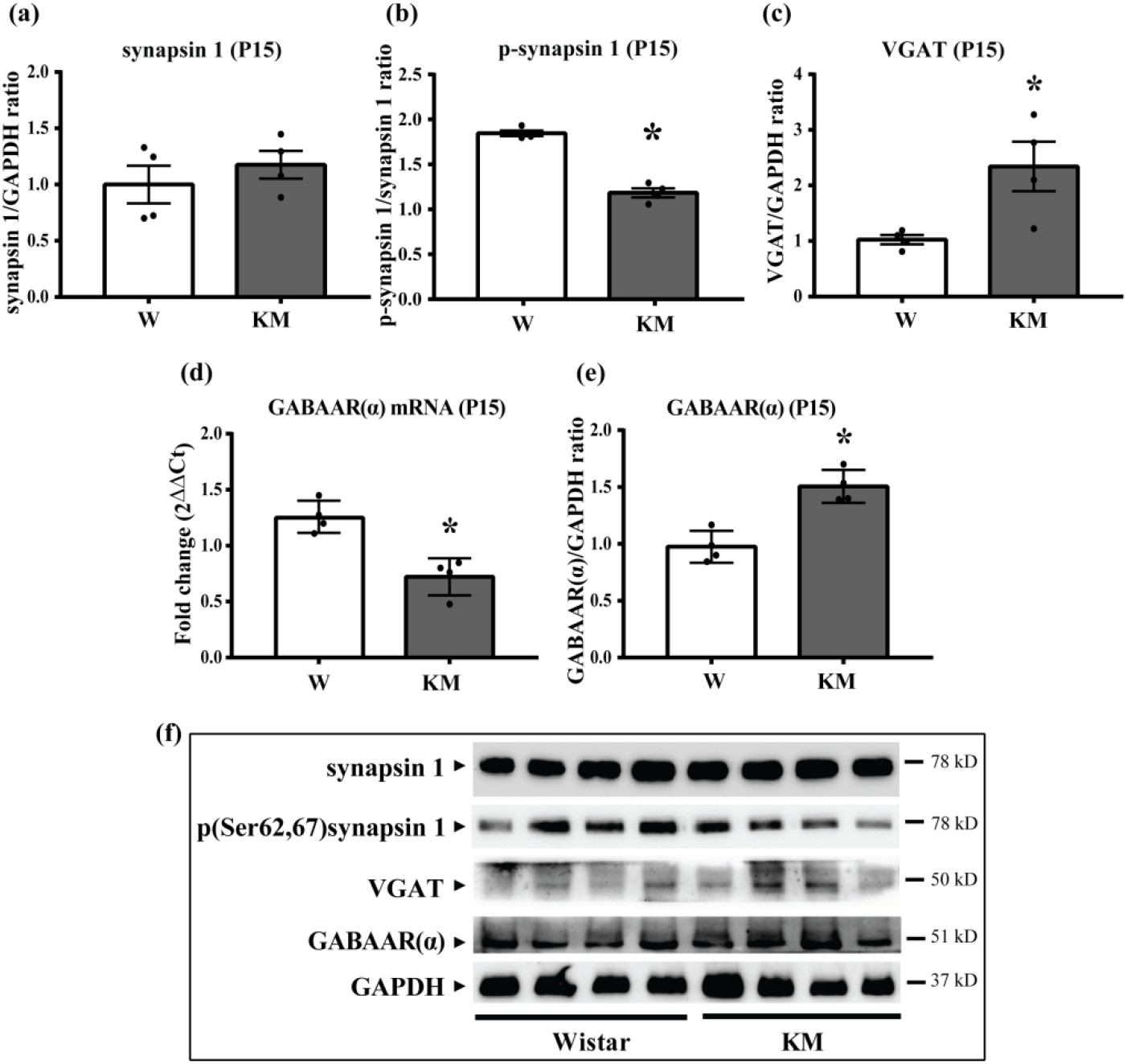
Analysis of pre- and postsynaptic proteins in the hippocampus of Wistar and KM rats at P15. **(a, b)** Synapsin 1 expression **(a)** in the hippocampus of KM rats did not differ from control Wistar (W) rats, but phosphorylated synapsin 1 (Ser62, 67) **(b)** was significantly lower. **(c)** VGAT expression was, in opposite, elevated. **(d, e)** Real-time PCR showed decrease in mRNA of GABAAR(α) **(d)** in the hippocampus of KM rats but corresponding protein content was higher than in Wistar control **(e)**. Data are presented as mean ± SEM. * – p<0.05 vs control. **(f)** Representative immunoblot images of synapsin 1, phospho(Ser62, 67)synapsin 1, VGAT, GABAAR(α), and glyceraldehyde 3-phosphate dehydrogenase (GAPDH) in the hippocampus of 15-day-old Wistar and KM rats.

Analysis of Cl^−^ transporters in the hippocampus was also performed. Expression of KCC2 and NKCC1 mRNA in the hippocampus of 15-days-old KM rats did not differ from the control (Figure 4a, b), however, protein expression of both KCC2 and NKCC1 evaluated by Western-blot analysis was significantly decreased (Figure 4c, e, f). Phosphorylation of KCC2 at Ser940, which is necessary for its stable binding with plasma membrane [36], was lower as well (Figure 4d, f). Immunohistochemical assay detected reduced KCC2 expression in the DG granular layer and CA4, while in the hilus it did not differ from the control (Figure 4g, i). Besides, at the 15^th^ day of postnatal development, KCC2 was practically absent in the inner part of the granular layer of both KM and Wistar rats. Probably, it was associated with gradual migration of new cells from the hilus to the granular layer during maturation of the hippocampus: cells which migrated earlier were detected in the outer part and already began expressing detectable KCC2 amount, while those who incorporated later were characterized with low KCC2 expression. In KM rats KCC2-immunopositive zone of the granular layer was significantly narrower than in the control (Figure 4h, i) indicating delayed KCC2 accumulation in the granular cells.

**Figure 4.**
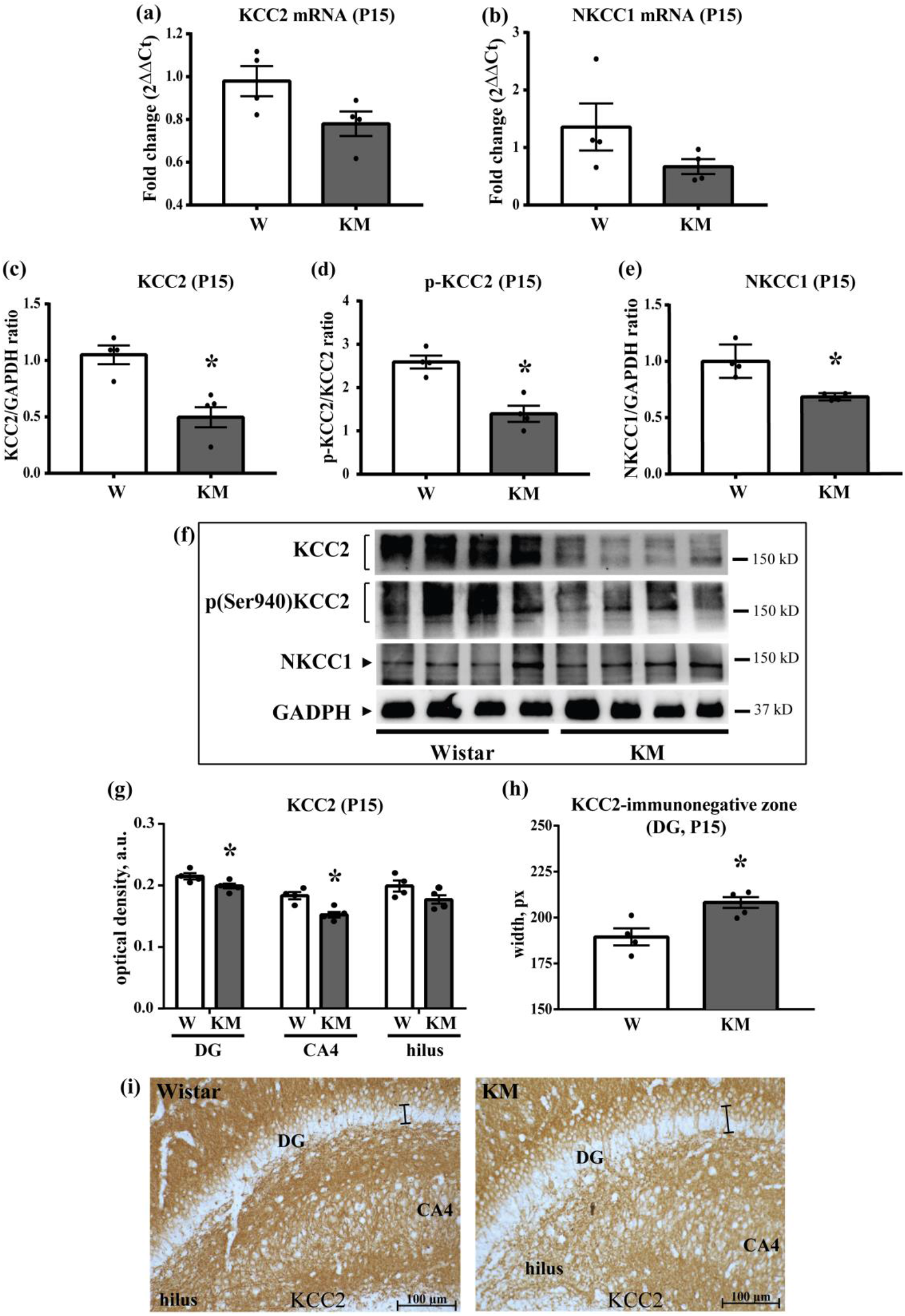
Analysis of chlorine transporters in the hippocampus of KM and Wistar rats at P15. **(a, b)** Both KCC2 **(a)** and NKCC1 **(b)** mRNA expression in the hippocampus did not differ between KM and Wistar (W) rats. **(c, d)** Expression of both KCC **(c)** and phospho(Ser940)KCC2 **(d)** estimated by Western blotting was reduced in the hippocampus of 15-days-old KM rats. **(e)** NKCC2 expression was as well lower than in the control. Data are presented in as mean ± SEM. * – p<0.05 vs control. **(f)** Representative immunoblot images of KCC2, phospho(Ser940)KCC2, NKCC1, and GAPDH in the hippocampus of Wistar and KM rats at P15. **(g)** Immunohistochemical analysis demonstrated significant decrease in KCC2 expression for the DG granular layer and CA4 of KM rats, while in the hilus expression was similar to Wistar control. Plot shows the optical density of immunopositive substance in arbitrary units (a.u.). **(h)** In the DG granular layer of KM rats, KCC2-immunonegative zone was broader than Wistar control. Plot demonstrates the width in pixels (px). Data are presented in as mean ± SEM. * – p<0.05 vs control. **(j)** Representative micrographs of KCC2 immunostaining in the hippocampus of KM and Wistar rats at P15. The width of KCC2-immunonegative zone is highlighted with the black line. Scale bars: 500 µm.

#### 60^th^ postnatal day (P60)

By the end of the second month of life (P60), KM rats demonstrated elevated numbers of both GAD67-positive (Figure 5a, c) and PV-positive (Figure 5d, f) neurons in all investigated hippocampal regions. At the same time, expression of both GAD67 (Figure 5b, c) and PV (Figure 5e, f) in the GABAergic fibers and terminals no longer differed between KM and Wistar rats. Observed alterations indicated elevated activity of GABAergic hippocampal neurons in KM rats.

**Figure 5.**
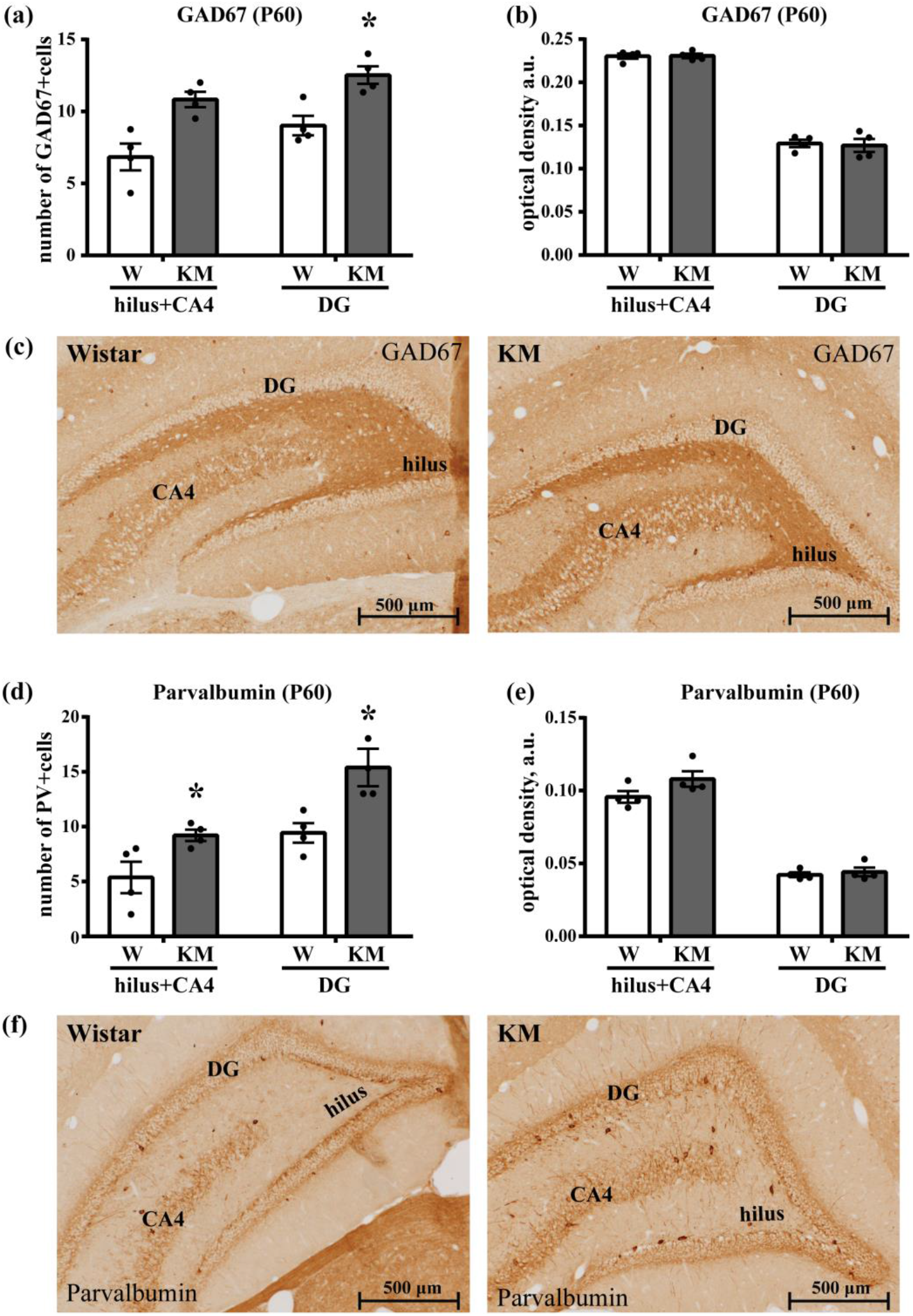
Expression of GAD67 and PV is upregulated in the hippocampus of KM rats at P60. **(a)** By the end of the second month of life, the number of GAD67-positive cells both in the hilus and CA4 zone and DG granular layer (DG) of KM rats was higher than in control Wistar (W) rats. **(b)** GAD67 expression in GABAergic fibers was similar to the control in both studied zones. **(c)** Representative micrographs of GAD67 immunostaining in the hippocampus of KM and Wistar rats at the age of two months. **(d, e)** Analysis of PV-positive cell populations **(d)** and PV expression in GABAergic fibers **(e)** showed similar manner of alterations. **(f)** Representative micrographs of parvalbumin immunostaining in the hippocampus of Wistar and KM rats at P60. Plots **(a)** and **(d)** demonstrate numbers of immunopositive cells; plots **(b)** and **(e)** – optical density of immunopositive substance in arbitrary units (a.u.). Data are presented as mean ± SEM. * – p < 0.05 vs control. Scale bars: 500 µm.

Analysis of markers responsible for synaptic transmission revealed a decrease in expression of synapsin 1 in the hippocampus of KM rats (Figure 6a, h), however, its phosphorylated form did not differ between KM and Wistar rats indicating no changes in synapsin activity (Figure 6b, h). In parallel, decrease in the hippocampal expression of both VGAT (Figure 6c, h) and GABAAR(α) (Figure 6d, h) was observed.

**Figure 6.**
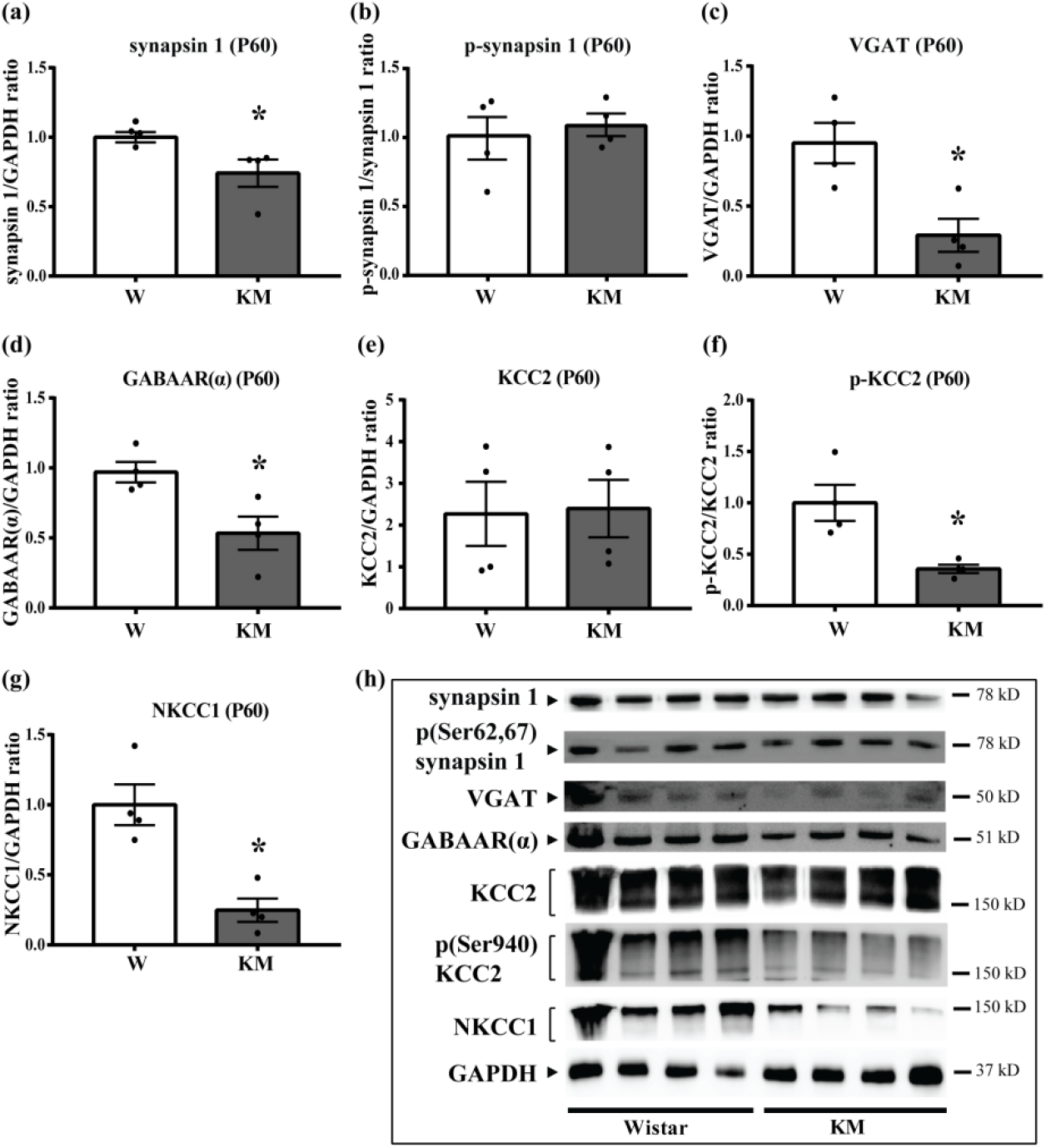
Analysis of pre- and postsynaptic proteins and chlorine transporters in the hippocampus of KM and Wistar rats at P60. **(a, b)** Two-month-old KM rats demonstrated decrease in total expression of synapsin 1 in the hippocampus **(a)**, while phospho(Ser62,67)synapsin 1 expression **(b)** was similar between KM and control Wistar (W) rats. **(c)** In the hippocampus of KM rats, significant decrease in VGAT expression was also observed. **(d)** Expression of GABAAR(α) in the hippocampus of KM rats was lower as well. **(e-g)** Total KCC2 expression **(e)** in the hippocampus of KM rats did not differ from Wistar control, however, both phospho(Ser940)KCC2 **(f)** and NKCC1 **(g)** expression was significantly lower. Data are expressed as mean ± SEM. * – p<0.05 vs control. **(h)** Representative immunoblot images of synapsin 1, phospho(Ser62,67)synapsin 1, VGAT, GABAAR(α), KCC2, phospho(Ser940)KCC2, NKCC1, and GAPDH in the hippocampus of KM and Wistar rats at P60.

In contrast to P15, two-months-old KM rats demonstrated unchanged hippocampal expression of KCC2 (Figure 6e, h), however, content of phosphorylated KCC2 (Figure 6f, h) and NKCC1 (Figure 6g, h) was still lower than in Wistar control.

#### 120^th^ postnatal day (P120)

In four-months-old KM rats, expression of GAD67 mRNA no longer differed from Wistar rats (Figure 7a). Nevertheless, populations of GAD67-positive cells and GAD67 expression in the GABAergic fibers were increased in all investigated hippocampal areas (Figure 7b-d). PV expression in GABAergic fibers also demonstrated ubiquitous increase (Figure 7f, g), and the numbers of PV-positive cells were elevated in the hilus and CA4, but not in the granular layer (Figure 7e, g).

**Figure 7.**
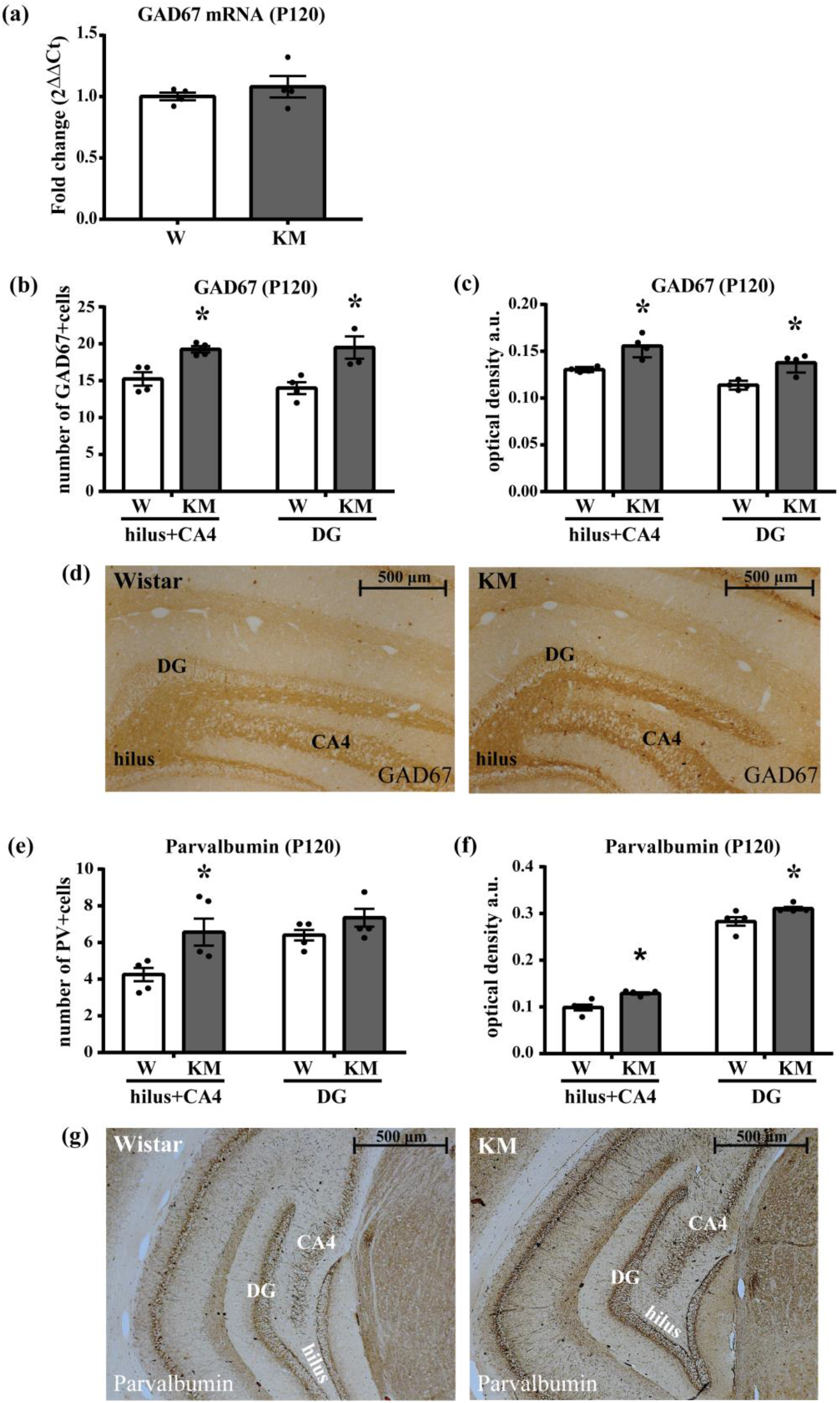
GAD67 and PV expression is still elevated in the hippocampus of KM rats at P120. **(a)** GAD67 mRNA did not differ between adult KM and Wistar (W) rats. Data are presented as mean ± SEM. * – p<0.05 vs control. **(b, c)** Both the numbers of GAD67-positive cells **(b)** and GAD67 expression in fibers **(c)** were ubiquitously increased in the hippocampus of KM rats. **(d)** Representative micrographs of GAD67 immunostaining in the hippocampus of Wistar and KM rats at P120. **(e)** KM rats demonstrated increased populations of PV-positive cells in the hilus and CA4 (hilus+CA4), but not in the granular layer (DG). **(f)** PV expression in GABAergic fibers was elevated in both studied hippocampal zones of KM rats. **(g)** Representative micrographs of PV expression in the hippocampus of Wistar and KM rats at P120. Plots **(b)** and **(e)** demonstrate the numbers of immunopositive cells, plots **(c)** and **(f)** – optical density of immunopositive substance in arbitrary units, a.u. Immunohistochemical data are presented as mean ± SEM. * – p<0.05 vs control. Scale bars: 500 µm.

These changes were accompanied with increased expression of total and phosphorylated synapsin 1 indicating upregulation of synaptic transmission in the hippocampus of KM rats (Figure 8a, b, f). VGAT expression (Figure 8c, f) and GABAAR(α) mRNA (Figure 8d) in four-months-old animals reached the control level. However, protein content of GABAAR(α) remained lower than in Wistar rats (Figure 8e, f).

**Figure 8.**
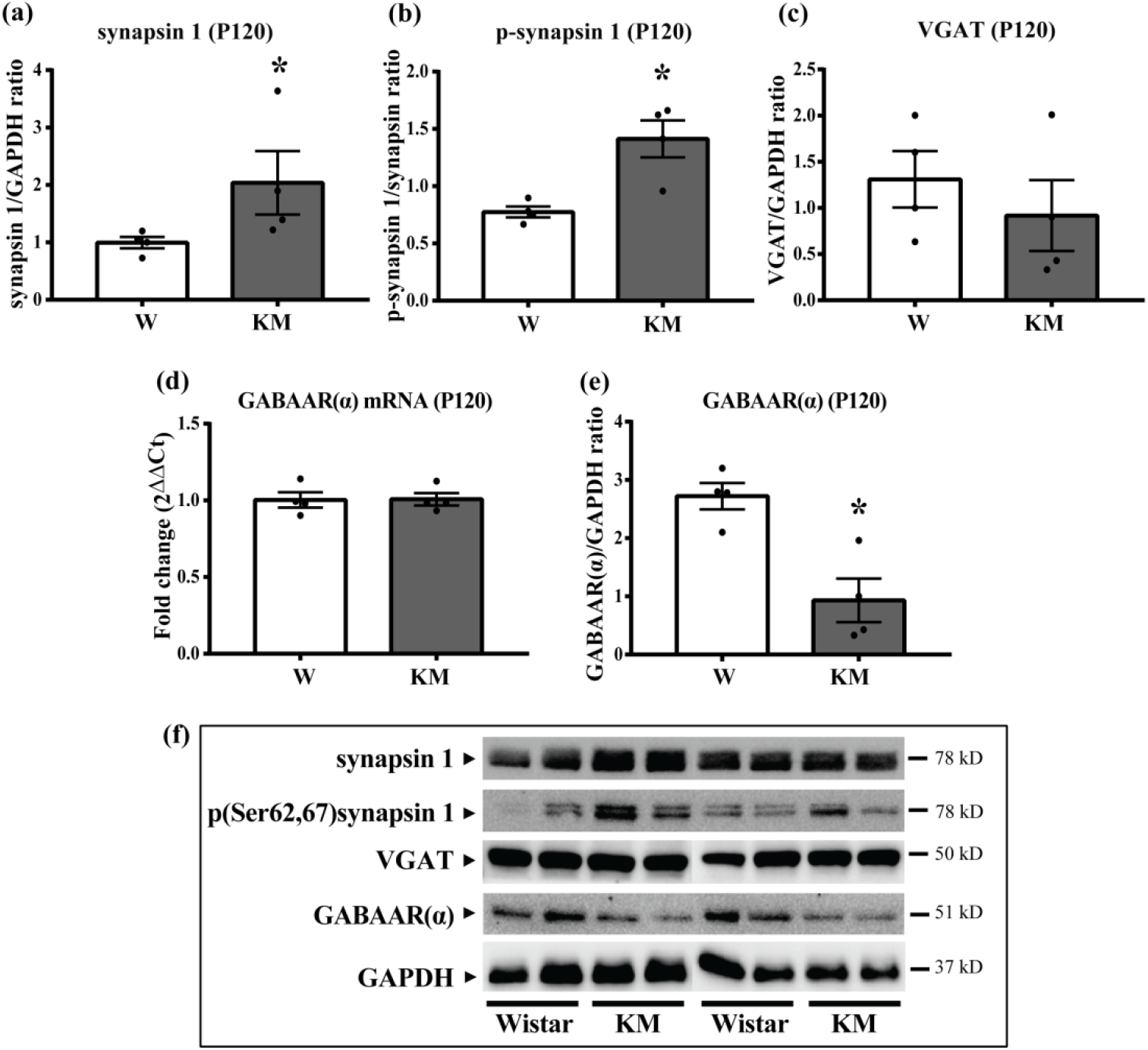
Analysis of pre- and postsynaptic proteins in the hippocampus of Wistar and KM rats at P120. **(a, b)** Expression of total **(a)** and phosphorylated **(b)** synapsin 1 was elevated in the hippocampus of four-months-old KM rats in comparison with Wistar control. **(c)** VGAT expression was similar in analyzed animal groups. **(d, e)** Evaluation of expression of GABAAR(α) in the hippocampus of KM rats revealed no changes in mRNA content **(d)**, but significant decrease in protein expression **(e)**. Data are presented as mean ± SEM. * p<0.05 vs control. **(f)** Representative immunoblot images of synapsin 1, phospho(Ser62,67)synapsin 1, VGAT, GABA-AR(α), and GAPDH in the hippocampus of Wistar and KM rats at P120.

Estimation of KCC2 expression revealed increased content of corresponding mRNA in the hippocampus of four-months-old KM rats (Figure 9a), however, expression of total and phosphorylated protein did not differ from Wistar control (Figure 9c-g). Immunohistochemical analysis also did not reveal any changes in KCC2 expression in studied hippocampal zones of KM rats, and, similar to Wistar rats, KCC2-immunonegative zone was no longer detected in the granular layer (Figure 9f, h). Both mRNA and protein expression of NKCC1 was unchanged as well (Figure 9b, e, g).

**Figure 9.**
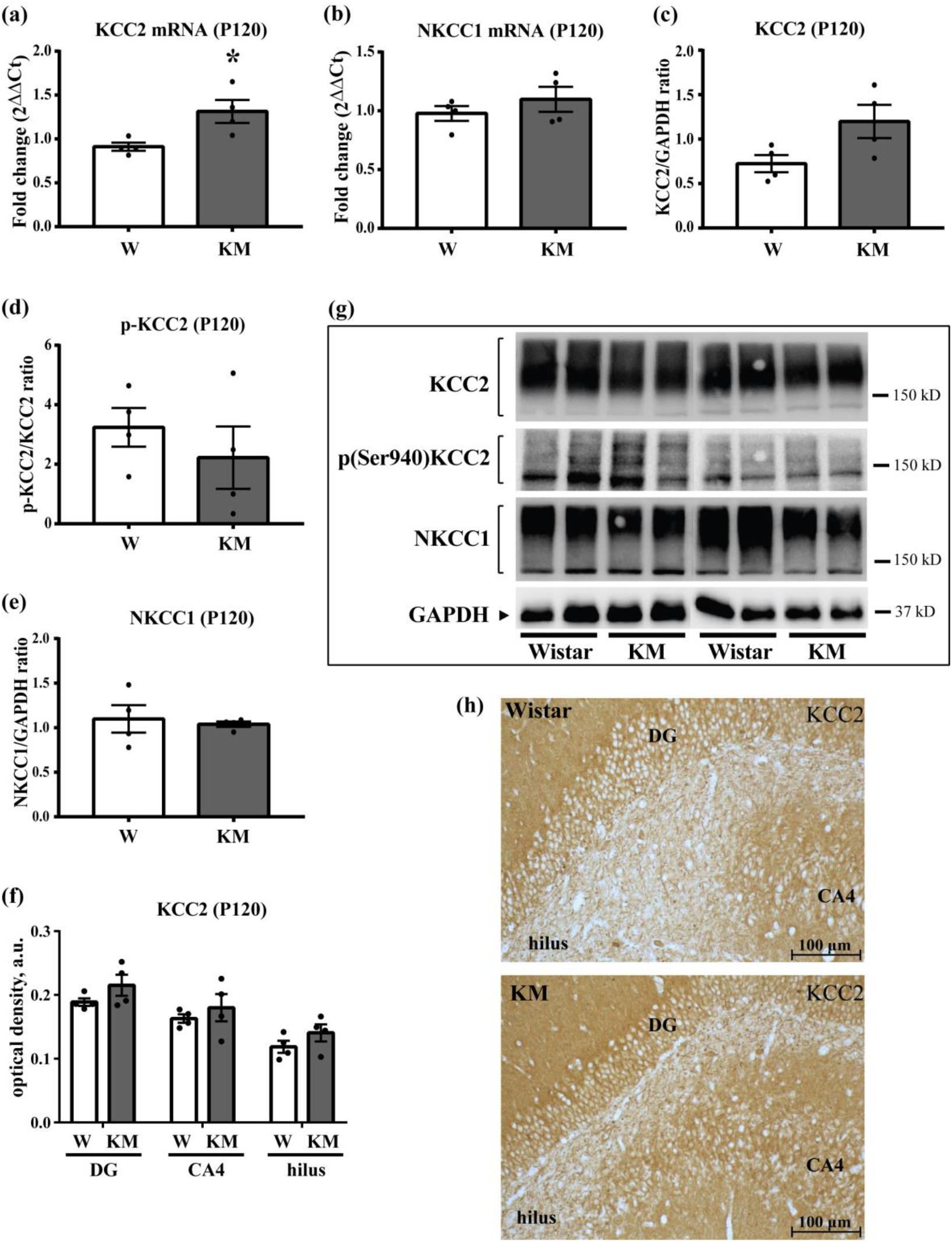
Analysis of chlorine transporters in the hippocampus of Wistar and KM rats at P120. **(a, b)** Real-time PCR revealed increase in KCC2 mRNA **(a)** in the hippocampus of KM rats as compared with Wistar (W) rats, but no change in NKCC1 mRNA **(b)** was observed. **(c-e)** Expression of KCC2 **(c)**, phospho(Ser940)KCC2 **(d)**, and NKCC1 **(e)** estimated with use of Western blotting demonstrated no alterations. Data are presented as mean ± SEM. * – p<0.05 vs control. **(f)** Representative immunoblots of KCC2, phospho(Ser940)KCC2, NKCC1, and GAPDH in the hippocampus of four-months-old Wistar and KM rats. **(g)** Immunohistochemical analysis also showed that KCC2 expression in DG granular layer, CA4, and hilus did not differ between Wistar and KM rats. Plot demonstrates optical density of immunopositive substance in arbitrary units, a.u. Immunohistochemical data are presented as mean ± SEM. * – p<0.05 vs control. **(h)** Representative micrographs of KCC2 immunostaining in the hippocampus of Wistar and KM rats at P120. Scale bars: 100 µm.

## Discussion

In the present work, we analyzed age-dependent changes in the expression and activity of key proteins responsible for production, synaptic transmission, and postsynaptic effects of GABA in the IC and hippocampus of KM rats at three ontogenetic points, P15, P60, and P120.

### 15^th^ postnatal day (P15)

According to the literature data, the end of the second postnatal week is a period when normal morphogenesis of both the IC and hippocampus is generally completed, although some developmental processes continue up to the end of the first month [21, 23, 37, 38]. Our previous data revealed that two-week-old KM rats still demonstrated increased proliferation and apoptosis in the IC and hippocampus indicating the delayed maturation of these structures in comparison to normal (Wistar) rats [27, 28]. Moreover, our previous studies showed that in the IC of KM rats at this time point the expression of GAD67 was similar to the control indicating no alterations in GABA synthesis, but the number of PV-positive cells was reduced pointing on decreased activity of GABAergic cells [29]. These alterations were accompanied with decreased activity of synapsin 1 suggesting decreased synaptic transmission and, in particular, attenuated GABA release [29]. Probably, these data pointed on the delayed formation of GABAergic system in the IC of KM rats at early steps of postnatal development.

Our present work provided new data complementing the picture of GABA dysregulation in the IC of KM rats during the first months of postnatal development. Thus, analysis of VGAT, responsible for GABA loading into synaptic vesicles, detected no changes in 15-day-old KM rats, but increased expression of GABA-A receptors pointed on reduced GABA binding and receptor internalization [39, 40] due to described earlier downregulation of GABAergic transmission.

Postsynaptic effects of GABA are tightly associated with Cl^−^ balance in the target cells which, in turn, depends on the expression and activity of chorine transporters KCC2 and NKCC1 [6]. Moreover, alterations in KCC2/NKCC1 expression are assumed to be the key mechanism that governs the switch of GABA effect from excitation to inhibition during postnatal development throughout the brain [41, 42]. At the first postnatal days, NKCC1 expression dominates, while KCC2 expression is low, supporting relatively high intracellular Cl^−^ and depolarizing GABA action [43, 44]. Then, during the first two weeks, active synaptogenesis is accompanied with an increase in KCC2 expression leading to enhanced Cl^−^ removal from the cells and establishment of hyperpolarizing GABA effect [24, 25]. Our study did not reveal any alterations in the expression of KCC2 in the IC of 15-day-old KM rats, however, abnormally increased expression of NKCC1 was observed. These data are insufficient to suppose any delay in excitatory-to-inhibitory GABA switch in the IC. Nevertheless, NKCC1 upregulation could be associated with increased Cl^−^ influx into GABA target cells leading to impaired inhibitory GABA action, in addition to decrease in GABA release.

Furthermore, our present study revealed impaired functional state of the GABAergic system in the hippocampus of 15-day-old KM rats. Thus, decreased GAD67 mRNA pointed on downregulation of this key enzyme responsible for GABA synthesis. In addition, abnormal distribution of GAD67 in somas and fibers indicated impaired trafficking of the protein into axon terminals of GABAergic cells. Expression of PV was lower as well suggesting decreased functional activity of GABAergic neurons.

In opposite, expression of VGAT in the hippocampus of KM rats was elevated at P15. Such changes might indicate increased GABA release into synaptic clefts, however, in parallel a decrease in activity of synapsin 1 was observed. According to published data, downregulation of synapsin 1 can be accompanied by increased VGAT content, but anyway it is associated with the inhibition of GABA release [45]. Thus, along with altered GAD67 and PV expression, changes of VGAT and synapsin 1 could indicate decreased GABA transmission in 15-day-old KM rats.

Expression of GABA-A receptors was also altered in the hippocampus of 15-day-old KM rats: thus, mRNA of the α1-subunit was decreased, while content of the corresponding protein, in opposite, was elevated. The literature data indicate that discordance between mRNA and protein expression is not rare: for different genes, changes in mRNA and protein profiles can demonstrate not only positive, but also negative or absent correlation [46]. In particular, it was shown that ischemic damage induced a decrease in mRNA of GABAAR(α) in the neocortex of rats, while protein expression was unchanged [47]. Similar alterations were observed for GABAAR(α) mRNA and protein in the cerebellum of aged rats [48]. However, the detailed mechanisms underlying such discrepancies are unknown. Possible explanations include different time scale of mRNA and protein synthesis and influence of other mechanisms responsible for mRNA and protein degradation or post-translational modifications [46]. In the present study, we supposed that increase in GABAAR(α) protein in the hippocampus of young KM rats, similar to the IC, could be associated with insufficient synaptic release of GABA resulting in its decreased binding and, in hence, decreased rate of receptor internalization, while the reason of decrease in corresponding mRNA remains to be elucidated.

Moreover, our results revealed down-regulation of both KCC2 and NKCC1 in the hippocampus of 15-day-old KM rats. Observed alterations could indicate an overall decrease in the activity of Cl^−^ transport that was accompanied with a decrease in GABA transmission. At the same time, the DG granular layer of KM rats contained wider KCC2-immunonegative zone, suggesting the excitatory GABA action in the greater amount of granular cells and postponed switch of GABA action from excitation to inhibition. Thus, our data pointed on considerable disturbance of chlorine balance in the hippocampal cells of KM rats.

Noteworthy, the relationship between KCC2 expression and postsynaptic GABA effect seems to be bidirectional. While there is a plenty of data describing KCC2-dependent switch of GABA effect, some studies have also documented that GABA action can potentiate KCC2 expression in cultured hippocampal neurons [49]. Thus, decreased KCC2 expression observed by us in the hippocampus of 15-day-old KM pups could also be the consequence of decreased GABA transmission.

Altogether, obtained data indicated that by the end of the second postnatal week, activity of GABAergic neurons and GABA transmission were abnormally decreased both in the hippocampus and the IC of KM rats. As this period is associated with active maturation of GABAergic system in these brain structures, our results suggest the genetically determined delay in the establishment of inhibitory GABAergic projections in the hippocampus and the IC of audiogenic KM rats.

Interestingly, when acting as excitatory neurotransmitter during the first days of postnatal development, GABA actively participates in the regulation of proliferation and differentiation of neural progenitors, migration of newborn neurons, and synaptogenesis [50, 51]. In particular, when exerting a depolarizing effect, GABA can stimulate migration of both glutamatergic and GABAergic neurons, while growth of KCC2 expression and decrease of intracellular Cl^−^ stops migration of cells [52]. Our previous results revealed impaired migration of newborn glutamatergic cells in the hippocampus of 15-day-old KM rats which led to abnormally high number of these cells in the hilus both in young and adult animals [28]. Thus, our present data let to suppose that these changes could be associated with prolonged excitatory action of GABA in a large number of the hippocampal cells during first two postnatal weeks.

### 60^th^ postnatal day (P60)

According to our previous investigations, by the end of the second postnatal month the development of both the IC and the hippocampus is completed in KM rats, as cell numbers, proliferation, and apoptosis no longer differed from Wistar control [27, 28]. However, elevated numbers of GAD67- and PV-positive cells and increased activity of synapsin 1 in the IC of two-month-old KM rats indicated upregulated activity of GABAergic system in this auditory center [29]. Our present data revealed also an increase in the expression of VGAT in the IC which additionally pointed on the higher activity of GABA release. Expression of GABA-A receptors and Cl^−^ transporters in the IC has reached the control level. Thus, as the IC is the critical structure responsible for triggering of AGS, observed enhancement of inhibitory transmission might be one of the main factors that prevent stable AGS expression in KM rats of this age.

In the hippocampus of KM rats, completion of morphogenesis was accompanied with the establishment of normal expression of GAD67 and PV in the terminals of GABAergic neurons, as well as with normalization of synapsin 1 activity. Moreover, similar to the IC, numbers of GAD67- and PV-positive neurons were elevated indicating an increase in functional activity of GABAergic cells.

Increased activity of GABAergic cells was accompanied with low expression of GABA-A receptors which could be explained by increased receptor internalization associated with enhanced GABA release. On the other hand, we also revealed a decrease in expression of VGAT, and simultaneous downregulation of both GABA receptors and vesicular transport of GABA could, in opposite, point on the attenuation of GABA transmission. In addition, down-regulation of Cl^−^ transporters, KCC2 and NKCC1, was observed, probably, indicating an overall decrease in the activity of Cl^−^ transport in GABA postsynaptic targets.

Thus, while in the IC of KM rats completed morphogenesis was associated with normalization and even upregulation of GABAergic system activity, in the hippocampus alterations were controversial: most of GABAergic markers showed normal or increased activity, while others indicated that some dysregulation still persisted.

### 120^th^ postnatal day (P120)

Analysis of adult KM rats with completely established ability of AGS expression also demonstrated considerable changes in the GABAergic system of the IC. Previously we detected abnormally low expression of GAD67 and PV in the IC central nuclei of adult KM rats indicating a decrease in functional activity of GABAergic neurons [29]. Our present data also revealed significant decrease in expression of KCC2 which let us to suppose a decrease in Cl^−^ efflux in GABA target cells and, in hence, impaired hyperpolarizing action of GABA in the IC. Thus, dramatic attenuation of GABAergic transmission, along with impaired Cl^−^ balance in GABA targets, might strongly contribute to the hyperexcitability of the IC in adult KM rats and mediate the onset of fully developed AGS.

In the hippocampus of adult KM rats, opposite changes in the GABAergic system were observed. Thus, considerable increase in both GAD67 and PV expression in the somas and processes of the hippocampal GABAergic neurons indicated upregulation of their functional activity and GABA production [34]. Increased activity of synapsin 1, together with normalization of VGAT expression and decreased content of GABA-A receptors, also confirmed activation of GABAergic transmission with increased receptor turnover. In addition, normalization of KCC2 and NKCC1 activity was observed indicating the restoration of Cl^−^ balance in GABA target cells, which is necessary for appropriate GABAergic inhibition.

Altogether, obtained data indicated significant increase in activity of GABA transmission in the hippocampus of adult KM rats. We suppose that such upregulation may provide a protective mechanism that prevents the involvement of the hippocampus in the spreading of epileptiform activity and, in hence, generalization of seizures at single sound stimulations.

Interestingly, the studies of other audiogenic rodents indicated opposite alterations in the GABAergic system of the IC and hippocampus. Thus, in the IC central nucleus of GEPR-9 rats, increased number of GAD67-positive neurons was observed indicating upregulation of GABA production [13]. In addition, genetic audiogenic seizure-prone hamsters from Salamanca (GASH:Sal) demonstrated unchanged expression of KCC in the IC and downregulation of the transporter in the hippocampus [20]. Probably, observed discrepancies can be explained by the innate variability of these audigenic animals: although AGS expression is similar, genetic basis (including the parental strain) and postnatal development of audiogenic sensitivity are different providing the basis for different neurochemical abnormalities.

## Conclusion

Results of the present study, together with previous findings [29], allowed us to compare postnatal development of GABAergic system in the IC and hippocampus of audiogenic KM rats and suggest a possible contribution of genetically determined alterations into AGS susceptibility. At early postnatal development (the end of the second postnatal week), KM rats demonstrated decreased activity of the key GABAergic markers in both the IC and hippocampus in comparison with Wistar rats. Additionally, altered expression of Cl^−^ transporters KCC2 and NKCC1 in both brain regions pointed on Cl^−^ imbalance in GABA target neurons and, in hence, impaired GABA-mediated inhibition. While in the IC up-regulation of NKCC1 was detected, in the hippocampus both transporters were decreased, and KCC2 was absent in abnormally wide area of the granular layer, indicating prolongation of excitatory GABA action. These results pointed on the delayed maturation of GABAergic system in both the IC and hippocampus that could underlie pathological changes further, in the adult KM rats.

By the end of the second postnatal month (P60), KM rats demonstrated stabilization and even up-regulation of GABA transmission in both the IC and hippocampus. However, in adult animals with fully developed readiness for AGS expression (P120), the opposite patterns of changes were observed in these brain structures. In the IC, functional activity of GABAergic cells was significantly decreased, while abnormally low KCC2 again let to suppose impaired GABA-mediated hyperpolarization in the IC neurons. On the other hand, previous analysis of excitatory (glutamatergic) system did not show any abnormalities in the IC of adult KM rats [29]. Thus, genetically determined dysfunction of inhibitory transmission might be one of the main factors of the IC hyperexcitability in KM rats, contributing to functioning of the IC as a “pacemaker” of AGS expression.

Conversely, in the hippocampus of adult KM rats, we revealed pronounced activation of GABAergic transmission, which was accompanied with normalization of Cl^−^ transporters activity. Noteworthy, our previous data demonstrated a decrease in activity of glutamatergic neurons in the hippocampus of adult naïve KM rats [53]. Thus, up-regulation of inhibitory transmission together with down-regulation of excitatory system can provide the mechanism that prevents the involvement of the hippocampus in the expression of single AGS.

## Material and Methods

### Animals

Male and female KM rats (Moscow State University, Russia) were used in our experiments. These rats were initially selected from Wistar rats and demonstrate clonic-tonic seizures in response to specific sound stimulation. Audiogenic susceptibility of KM rats develops during postnatal ontogenesis being fully formed at the age of 3 – 3.5 months.

To reveal genetically determined aberrations in GABAergic system of the IC and the hippocampus, we recruited naïve KM rats which were not previously exposed to sound stimulation and had no experience of AGS expression prior to sacrificing. Three age groups of KM rats were investigated: 1) animals at the age of 15 days (P15, n=8) when active morphogenesis of the IC and the hippocampus occurs; 2) two-month-old animals (P60, n=8) with completed development of both structures of interest; 3) four-months-old animals (P120, n=8) which demonstrate fully developed ability to express of AGS. Wistar rats of corresponding ages were used as a control (n=8 for each age group). All animals were housed in standard vivarium cages under 12/12 light-dark cycle with *ad libitum* access to food and water.

### Sample preparation

Four animals from each experimental group were deeply anesthetized with i.p. injection of Zoletil/Xylazine mixture (60 mg/kg+10 mg/kg; Virbac, France), perfused transcardially with phosphate buffered saline (PBS) followed by 4% paraformaldehyde (PFA) and decapitated. The brains from all animals were removed, postfixed in 4% PFA at +4 °C (5 days), subsequently immersed in 15% sucrose for cryoprotection (3 days), then frozen, and stored at -80°C for further immunohistochemical analysis.

The other four animals from each group were decapitated; brains were removed, divided into hemispheres, and then the IC and dorsal hippocampi were dissected. The IC from both hemispheres of all these animals were used for Western blot analysis. In two-months-old Wistar and KM rats, both hippocampi were collected only for Western blotting. For each animal from P15 and P120 groups, the hippocampus from one hemisphere was used for Western blot analysis and from the other – for real-time PCR.

### Immunohistochemistry

The frozen brains were sectioned coronally at 10 µm using Leica cryostat. Immunohistochemical analysis was carried out according to standard biotin-streptavidin protocol. Cut cryosections (10 μm) containing the hippocampus or IC were incubated with primary antibodies (Table 1) overnight at room temperature. Thereafter sections were washed in PBS, incubated for 1 h with biotinylated secondary antibodies (Table 1) followed by incubation with рeroxidase-streptavidin complex (1:500, Supelco, #S2438) for 1 h. The peroxidase reaction was revealed in the buffer containing 3,3′-diaminobenzidine (DAB, Sigma-Aldrich, #D5637) and hydrogen peroxide (0.01%). To examine the specificity of staining, we performed the negative control (the same protocol without primary antibodies) that demonstrated no staining. Finally, the sections were dehydrated and coverslipped.

**Table 1.** Used antibodies for immunohistochemistry (IHC) and Western blotting (WB).

| Antibody | Manufacturer | Dilution for IHC | Dilution for WB |
| --- | --- | --- | --- |
| **Primary antibodies** |  |  |  |
| KCC2 Polyclonal Antibody | Invitrogen, #PA5-78544 | 1:250 | 1:1000 |
| Na-K-Cl cotransporter Monoclonal Antibody | DSHB Hybridoma Product, # t4 | 1:100 | 1:500 |
| GABA-AR alpha 1 Antibody | Novusbio, #NB300-191 | 1:100 | 1:1000 |
| Anti-GAD67 Antibody, clone 1G10.2 | Millipore, #MAB5406 | 1:500 | - |
| Parvalbumin Polyclonal Antibody | TermoFischer scientific #PA1-933 | 1:100 | - |
| Synapsin I Antibody | Novusbio, #NB300-104 | - | 1:1000 |
| Synapsin I [p Ser62, p Ser67] Antibody | Novusbio, #NB300-745 | - | 1:1000 |
| VGAT Polyclonal Antibody | Invitrogen, #PA-27569 | - | 1:1000 |
| GAPDH (14C10) Rabbit mAb | Cell Signaling Technology, #2118 | - | 1:1000 |
| **Secondary antibodies** |  |  |  |
| Goat Anti-Mouse IgG antibody (H+L), biotinylated | Vector Laboratories, #BA-9200 | 1:500 | - |
| Goat anti-rabbit IgG antibody (H+L), biotinylated | Vector Laboratories, #BA-1000 | 1:500 | - |
| Anti-rabbit IgG (whole molecule)–peroxidase antibody produced in goat | Sigma-Aldrich, #A0545 | - | 1:10000 |
| Anti-Mouse IgG (whole molecule)–Peroxidase antibody produced in rabbit | Sigma-Aldrich, #A9044 | - | 1:40000 |

### Evaluation of sections

The sections were processed under standardized conditions in every experiment, i.e. control and experimental groups in each experiment were collected, fixed, and processed for analysis simultaneously. Analysis of immunostaining in the IC and dorsal hippocampus was performed with use of images taken by a 20x/0.5 objective on Zeiss Axio Imager A1 microscope (Carl Zeiss Microscopy GmbH). Every 15th section was selected for analysis of each investigated protein. In total, five sections of the studied zone per each immunostaining were analyzed for each animal.

Expression of GABAAR(α), KCC2, and NKCC1 in the IC central nucleus, as well as expression of GAD67, PV, and KCC2 in the dentate gyrus (DG) granular layer, hilus, and CA4 subfield of the dorsal hippocampus, were estimated as optical density of immunopositive substance on 8-bit images with use of ImageJ software (version 6.0). Optical density of the background was estimated at the same slice in non-immunoreactive brain tissue. Numbers of GAD67- and PV-positive cells in the same hippocampal areas were also counted. In P15 animals, we additionally analyzed the width of KCC2-immunonegative zone of the granular layer. The width was calculated for each hippocampus at the same level in the medial-lateral direction by ImageJ and then averaged for each animal.

### Western blot analysis

Hippocampal and IC samples were homogenized in lysis buffer (20 mM Tris pH 7.5, 1% Triton-X100, 100 mM NaCl, 1 mM EDTA, 1 mM EGTA) containing protease inhibitors (Sigma-Aldrich, #P8340) and phosphatase inhibitor cocktail (Roche, #04 906 837 001) using tissue grinder at 4°C. Insoluble materials were removed by centrifugation. Total protein content in samples was determined by Lowry assay with bovine serum albumin (BSA) as a standard. The supernatant was mixed in ratio 2:1 with 3x loading buffer (0.2 M Tris-HCl pH 6.7, 6% sodium dodecyl sulfate, 15% glycerol, 0.003% bromophenol blue, and 10% β-mercaptoethanol) and incubated for 10 min at 96°C. Equal amounts of samples (10 μg protein per line) were loaded for electrophoresis, proteins were separated on 10% polyacrylamide gel, and then transferred to a nitrocellulose membrane (Santa Cruz Biotechnology, #sc-3718). The membranes were incubated in 5% non-fat milk or 3% BSA in Tris buffered saline with Tween (TBST) buffer (0.1% Tween 20, 20-mM Tris, 137-mM NaCl; pH 7.4) for 1 h and then incubated overnight with primary antibodies (Table 1). Thereafter the membranes were washed in TBST buffer and incubated with peroxidase-conjugated secondary anti-rabbit or anti-mouse antibodies (Table 1) for 1 h at room temperature. Specific protein bands were visualized by chemiluminescent reaction produced by SuperSignal™ West Dura Extended Duration Substrate (ThermoFisher Scientific, #34075) with use of ChemiDoc MP Imaging System (#12003154, Bio-Rad Laboratories Inc., Hercules, CA, USA).

Densitometric analysis of protein content was performed using ImageJ software (version 6.0). Expression of VGAT in the IC was estimated by normalizing to actin. Expression of phospho(Ser940)KCC2, KCC2, NKCC1, phospho(Ser62,67)synapsin 1, synapsin 1, GABAAR(α), and VGAT in the hippocampus was estimated by normalizing to GAPDH. The hippocampal expression of phospho(Ser940)KCC2 and phospho(Ser62,67)synapsin 1 was evaluated by normalizing to the total form of the corresponding protein. Detection of proteins used for normalization was performed with use of the same membranes, as detection of proteins of interest.

### cDNA synthesis and real-time PCR

Total RNA was isolated from hippocampal samples using PureZOL™ RNA isolation reagent (BioRad, #732-6890). RNA concentration was measured using a CLARIOstar Plus spectrophotometer (BMG LABTECH) following the standard procedure. The purity of RNA samples was verified by confirming an optical density ratio A260/A280>1.8.

Two μg of total RNA was used for cDNA synthesis using RevertAid First Strand cDNA Synthesis Kit (Thermo Scientific, #K1622). Quantitative real time RT–PCR was performed using Evrogen 5x qPCR mix-HS SYBR (Evrogen, #PK147L). Primers (Table 2) were designed with the Primer-BLAST software (NCBI, USA).

**Table 2.** Primers for qPCR

| Name | 5'-3' sequence |
| --- | --- |
| NKCC1 | F: tcg-tga-gag-gag-gag-gag-cat-ac |
|  | R: aga-tgc-cca-gaa-gaa-cca-cca-c |
| KCC2 | F: gca-gag-gtg-gaa-gtc-gtg-gag |
|  | R: ttc-cgg-ctc-gtt-ctt-ggt-g |
| GABA-A<br>receptor α-<br>subunit | F: agc-ctg-cat-tta-aga-aca-ga |
|  | R: gca-aca-gtg-aag-tta-tga-gc |
| GAD67 | F: gct-gga-agg-cat-gga-agg-ttt-ta |
|  | R: acg-ggt-gca-att-tca-tat-gtg-aac-ata |
| GAPDH | F: tcc-ctc-aag-att-gtc-agc-aa |
|  | R: aga-tcc-aca-acg-gat-aca-tt |

Real-time PCR analysis was performed with use of LightCycler 96 real-time PCR system (Roche). The PCR parameters were following: initial denaturation - one cycle at 95 °C for 15 min; 40 cycles of denaturation, amplification, and quantification (95 °C for 15 s, 60 °C for 30 s, and 72 °C for 5 s); the melting curve - start at 65 °C and gradual increase to 95 °C. Cycle thresholds were normalized against the reference gene GAPDH. Expression in control hippocampi was estimated as “1”. Relative fold expression of genes was calculated in Microsoft Excel by the 2-ΔΔCt method. The results are given as bar charts. Each value was combined from 3 independent PCR replicates for each cDNA sample.

### Statistical analysis

All data obtained with use of Western blot, real-time PCR, and immunohistochemical analyses were processed statistically by the Mann–Whitney U test using GraphPad 7 Software. Results are presented as mean ± standard error of mean (SEM). Differences were regarded as significant at p<0.05.

## Author Contributions

Conceptualization, A.A.N., E.V.C.; methodology, A.A.N., E.V.C.; formal analysis, A.P.I., S.D.N.; investigation, A.P.I., S.D.N., A.A.N.; data curation, E.V.C.; writing—original draft preparation, A.P.I., E.V.C.; writing—review and editing, A.A.N., M.V.G.; visualization, A.P.I., S.D.N., A.A.K.; supervision, M.V.G.; funding acquisition, A.A.N. All authors have read and agreed to the published version of the manuscript.

## Funding

This study was supported by the Russian Science Foundation (RSF) Grant № 22-75-00060, https://rscf.ru/project/22-75-00060/.

## Institutional Review Board Statement

All experiments with rats were conducted in accordance with EC Directive 86/609/EEC for animal experiments and approved by the Institutional Animal Care and Use Committee at the Sechenov Institute of Evolutionary Physiology and Biochemistry. (protocol code 6-3/2022, June 23, 2022).”

## Data Availability Statement

The data are available on a reasonable request to the corresponding author.

## Acknowledgments

Part of the analysis was done at Research Resource Center #441590 at Sechenov Institute of Evolutionary Physiology and Biochemistry, the Russian Academy of Sciences.

## Conflicts of Interest

The authors declare no conflict of interest.

## Notes

### Competing Interest Statement

The authors have declared no competing interest.

